# Is methylation of the CTLA-4 gene promoter region an epigenetic mechanism of autoimmune thyroiditis in hepatitis C? *In silico* experimental observation

**DOI:** 10.1101/2020.07.17.209650

**Authors:** Gabriela Correia Matos de Oliveira, Aline Zamira Freire Teles Aragão, Luís Jesuíno de Oliveira Andrade

## Abstract

**Introduction:** Cytotoxic T lymphocyte-associated antigen 4 (CTLA-4) is a crucial immune control point receptor that regulates T cell activation. Epigenetic mechanisms, such as DNA methylation and histone modifications, modulate DNA packaging in the nucleus and influence Gene expression. Autoimmune thyroiditis may be associated with hepatitis C virus (HCV) infection as well as the CTLA-4 Gene.

**Objective:** To in silico simulate the methylation of the promoter region of CTLA-4 gene as an epigenetic factor triggering autoimmune thyroiditis by HCV.

**Methods:** We analyzed by in silico simulation the hypermethylation scenarios of the CTLA-4 Gene promoter region, aligning CTLA-4 and HCV sequences (genotypes 1, 2 and 3) through BLAST software - http://blast.ncbi.nlm.nih.gov/Blast.cgi, and identifying their methylated and unmethylated CpG sites. After the sequences obtained with the alignment of the methylation points by MultAlin program, the consensus sequences obtained were submitted to the BLAST similarity search. The GC content calculation and HCV annotation were performed using ENDMEMO (http://www.endmemo.com/bio/gc.php). The MethPrimer was used to identify and locate the methylation CpGi within the HCV genome.

**Results:** The location of CTLA-4 on chromosome 2 and the alignment of the amino acid sequences are presented: CTLA-4 and HCV genotype 1, CTLA-4 and HCV genotype 2 and CTLA-4 and HCV genotype 3 are presented, as well as the methylation sites.

**Conclusion:** In susceptible individuals, hypermethylation promotes reduced CTLA-4 expression and increases the risk of autoimmune thyroiditis in HCV-infected individuals.

## Introduction

Cytotoxic T lymphocyte-associated antigen 4 (CTLA-4) a member of the immunoglobulin superfamily, is a crucial immune control point receptor that regulates T cell activation, has been identified as a risk factor for certain autoimmune diseases, becoming recognized as a risk agent for various T-cell-mediated autoimmune diseases (1,2).

The conception of epigenetics was first time suggested by Conrad Waddington in 1942 (3). Epigenetic occurrences lead to heritable alterations in gene manifestation another than the changes in DNA nucleotide sequences (4). The DNA-methylation is the most easily evaluated, and likely the most stead epigenetic characteristic, with important regulatory functions in animals. Epigenetic mechanisms, such as histone modifications or incorporation of variant histones, and DNA-methylation regulate DNA packaging in nucleus and act in gene expression, being capable design your own heredity via methylation of hemimethylated sites after DNA replication by preservation DNA-methylases (5,6). Histone transformations could be associated to reader-protein connection complexes to regulate gene expression, and favor to epigenetic heritance or to revert epigenetic marks in the chromatin (7). The histone deacetylation, DNA-hypermethylation or DNA-hypomethylation, and microRNAs in control of Gene expression are epigenetic changes that can reciprocal action (8).

The interactions of complex virus-host-environment demonstrate that epigenetic mechanisms occur soon after hepatitis C virus (HCV) infection, once a new epigenetic condition recorded and eternalized in later mitosis, originating in the reprograming of the cell transcription (9).

A T-cell response is essential for response of HCV infection, and co-stimulatory molecules decrease T-lymphocyte responses by connection with CTLA-4 (10). The epigenetic mechanism of DNA-methylation in CTLA-4 Gene in autoimmune thyroiditis happens in region of 5’promoter regions with high density and principally results in transcriptional suppression, and this DNA-methylation has results a fault to establishment and maintenance immunologic non-responsiveness or tolerance to self-antigens (11). Thus, autoimmune thyroiditis may be associated with HCV infection as well as the CTLA-4 Gene.

The ease of the availability of vast amounts of sequence data, associate to advances in computational biology facilitates *in sílico* analysis of several molecular framework. The aim of our study was to *in sílico* simulate the methylation of the promoter region of CTLA-4 Gene as an epigenetic factor triggering autoimmune thyroiditis by HCV infection.

## Method

We analyzed by *in sílico* simulation the methylation scenarios of the CTLA-4 Gene promoter region, aligning the CTLA-4 and HCV sequences (genotypes 1, 2 and 3), and identifying the methylation of genomic regions known as CpG islands (CpGi) sites.

### Sequences analysis

We got the CTLA-4 and HCV sequences (genotypes 1, 2 and 3) through the FASTA format in the NCBI database (http://www.ncbi.nlm.nih.gov/), with the aim of identifying possible regions which methylations.

A sample of thyroid peroxidase [Homo sapiens] was acquired at NCBI database Accession: AAA61217.2 GI: 4680721.

The following HCV sequences were obtained: polyprotein [Hepatitis C virus genotype 1], Accession: NP_671491.1 GI: 22129793; polyprotein [Hepatitis C virus genotype 2], Accession: YP_001469630.1 GI: 157781213; and polyprotein [Hepatitis C virus genotype 3], Accession: YP_001469631.1 GI: 157781217.

A sample of CTLA-4 [Homo sapiens] was acquired at GenBank Accession: AAL07473.1 GI: 15778586.

After the sequences acquired with the alignment of the methylation points by MultAlin program, the consensus sequences acquired were submitted to the BLAST software similarity evaluation. MultAlin is a digital platform supported on an algorithm exploiting gradual paired alignment considering the relationships that may exist between some subsets of the sequels. The application add choices to group the intended output model and its dimension and colors besides configure individualized alignment specifications like the quantity of repetitions and gap punishment obligatory.

The guanine-cytosine (GC) content calculation and HCV annotation were performed using ENDMEMO (http://www.endmemo.com/bio/gc.php).

Primer nucleotide sequence of CTLA-4 was verified using Primer-BLAST in NCBI web site.

### Analysis of methylation

The MethPrimer Express^®^ Software v1.0 (Applied Biosystems) a web-based program which has been developed in 2002 by Li & Dahiya (12) was used to identify and locate the methylation CpGi within the HCV genome. MethPrimer is an online platform which provides a number of tools and databases to become possible the investigation of epigenetics and of DNA methylation, incorporating tools to project probes and primers for several bisulfite conversion based PCRs, predicting CpGi, and manipulating sequences.

## Results

The NCBI show that CTLA-4 has cytogenetic location in the 2q33.2, where is the long (q) arm of chromosome 2 at situation 33.2, and molecular localization in the base pairs 203,867,771 to 203,873,965 on chromosome 2 (Figure 1).

**Figure 1.**
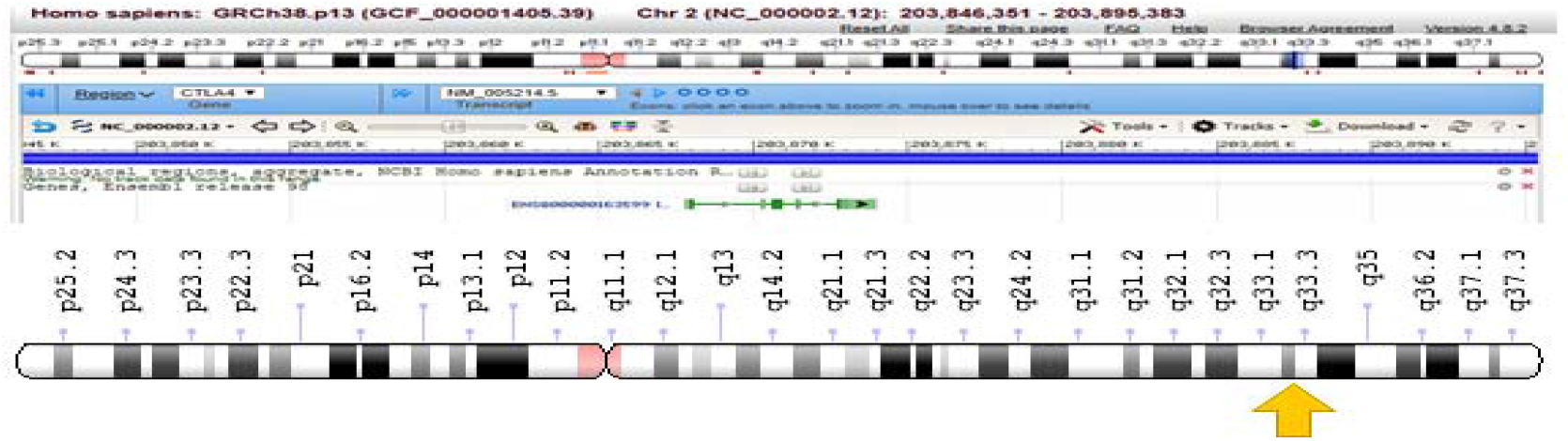
Location of CTLA-4 on chromosome 2 (Source NCBI).

### Identification of CpGi

The default parameters as proposed by Deaton and Bird (13) were used to identify the potential CpGi. The parameters were as follows: length of DNA sequence > 300bp; CpG checked/CpG expected proportion > 0.6 and C+G% > 50%.

### Thyroid peroxidase [Homo sapiens] - CpGi Prediction

The *in silico* analysis and the primers for quantitative DNA methylation analysis of thryroid peroxidase, and the sequence alignment of CTLA-4 and Thyroid peroxidase [Homo sapiens] not revealed the presence of putative promoters and none CpGi (Figure 2).

**Figure 2.**
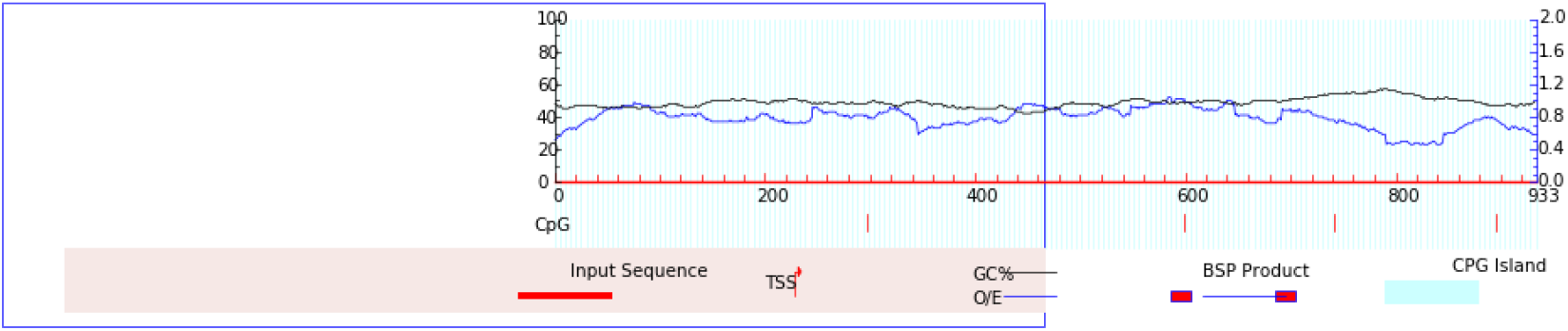
Prediction of methylation sites in the CpGi from Thyroid peroxidase [Homo sapiens].

### Hepatitis C virus genotype 1, complete genome - CpGi Prediction

The *in silico* analysis and the primers for quantitative DNA methylation analysis of HCV genotype 1 Gene, and the sequence alignment of the HCV genome and CTLA-4 demonstrated the existence of two putative promoters and two CpGi in the 5’ genomic region (Figure 3 & 4).

**Figure 3.**
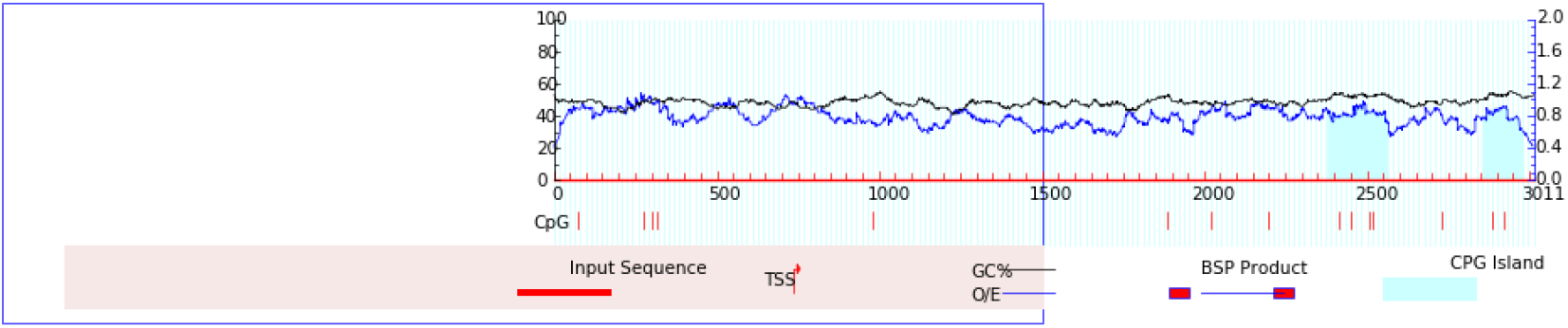
Prediction of methylation sites in the CpGi from Hepatitis C virus genotype 1.

**Figure 4.**
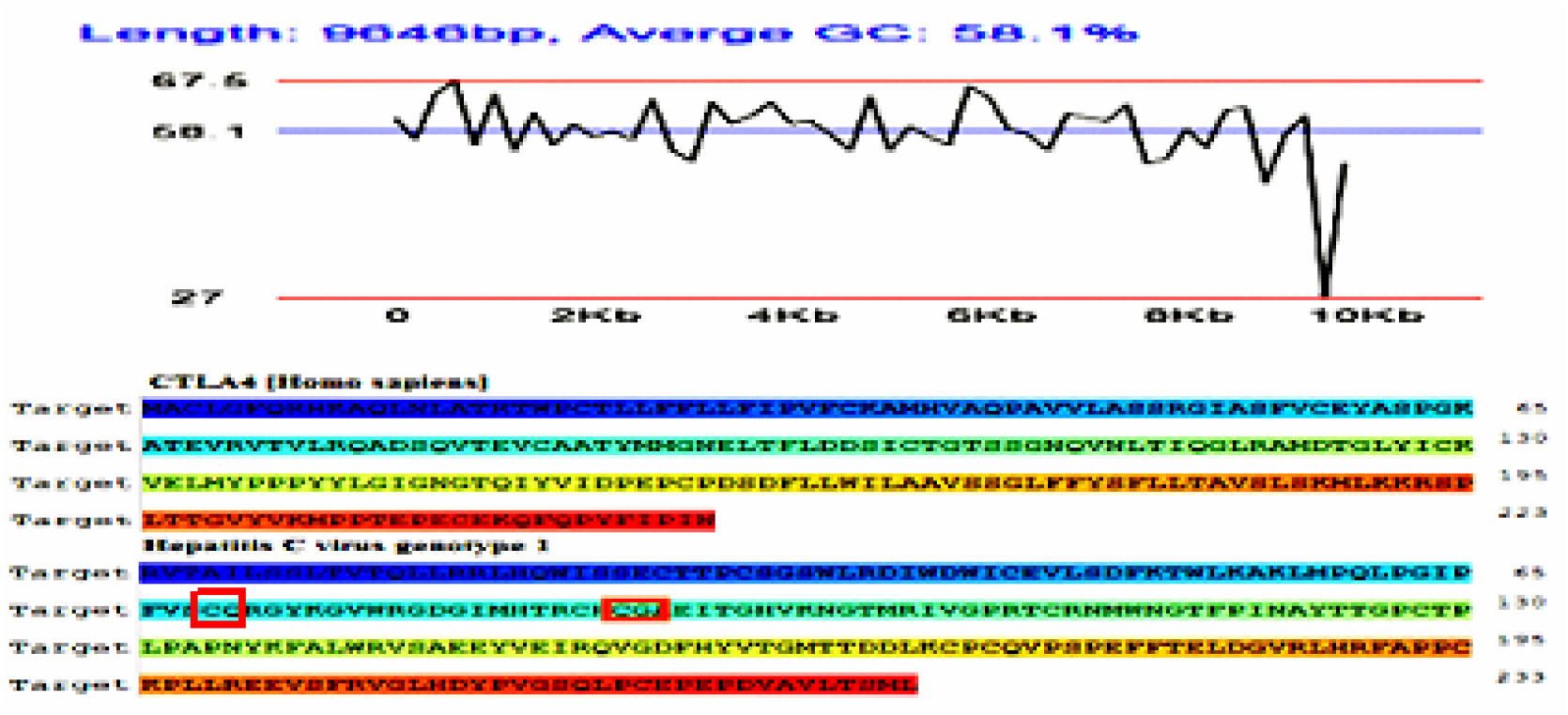
GC Content Distribution of HCV genotype 1.

### Hepatitis C virus genotype 2, complete genome - CpGi Prediction

The parameters analysis and the primers for quantitative DNA methylation analysis of HCV genotype 2 Gene, and the sequence alignment of CTLA-4 and Hepatitis C genome revealed the presence of one putative promoters and one CpGi in the 3’ genomic region (Figure 5 & 6).

**Figure 5.**
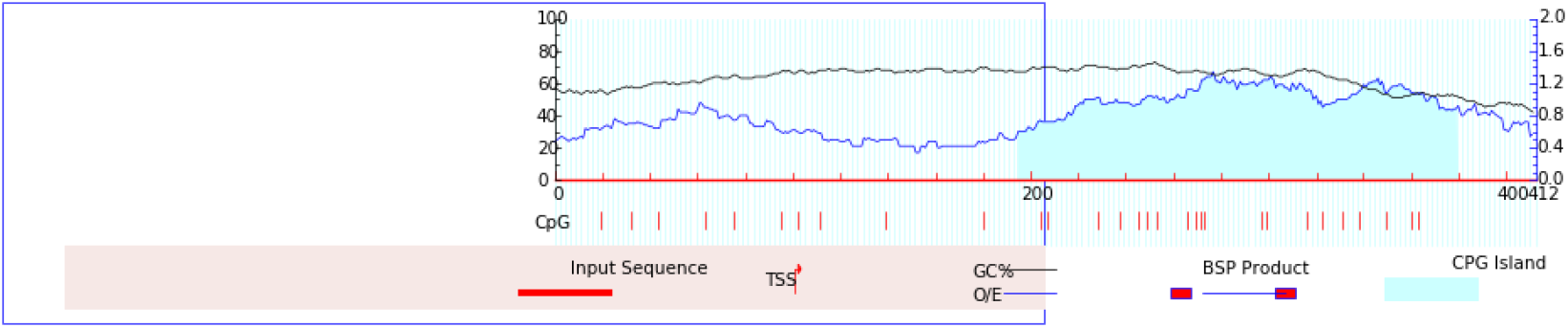
Prediction of methylation sites in the CpGi from HCV genotype 2.

**Figure 6.**
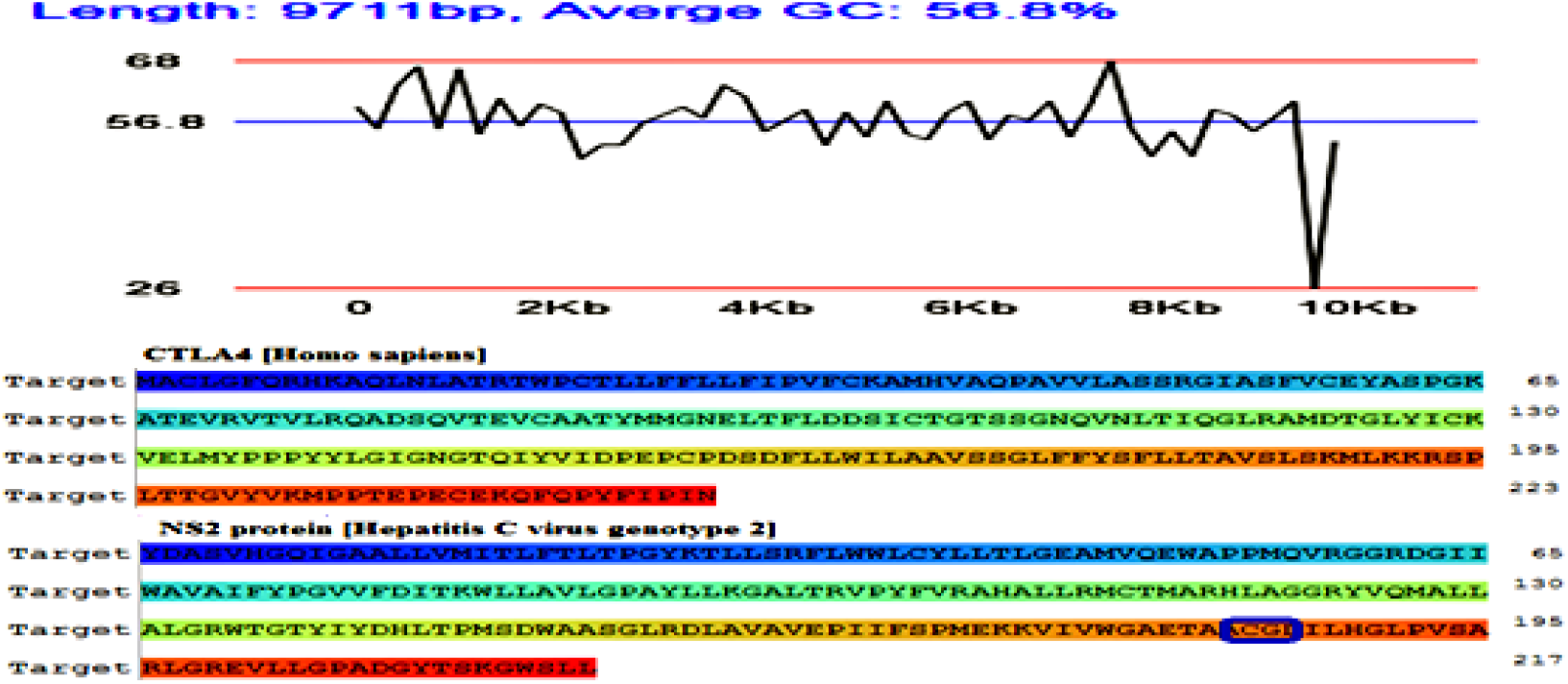
GC Content Distribution of HCV genotype 2.

### Hepatitis C virus genotype 3, complete genome - CpGi Prediction

The *in silico* analysis and the primers for quantitative DNA methylation analysis of HCV genotype 3 Gene, and the sequence alignment of CTLA-4 and Hepatitis C genome revealed the presence of two putative promoters and two CpGi in the 5’ genomic region (Figure 7 & 8).

**Figure 7.**
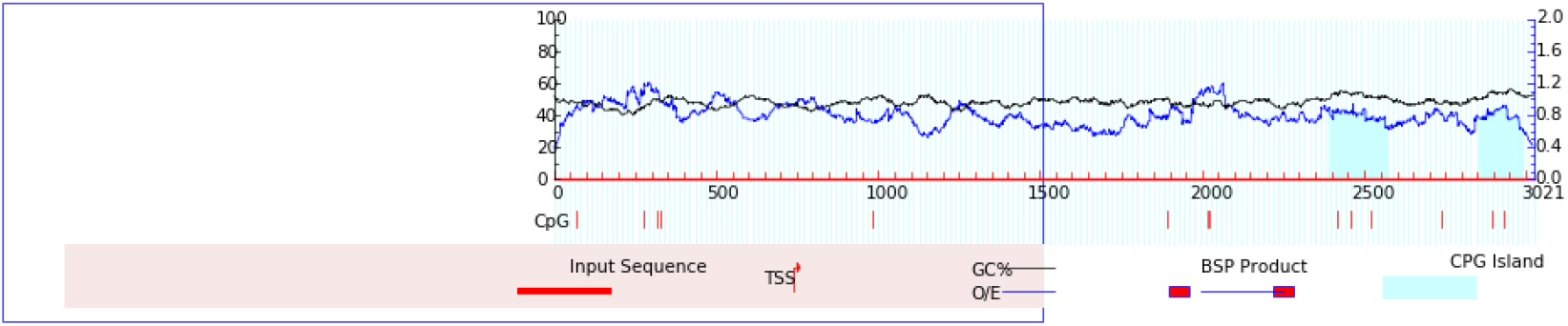
Prediction of methylation sites in the CpGi from HCV genotype 3.

**Figure 8.**
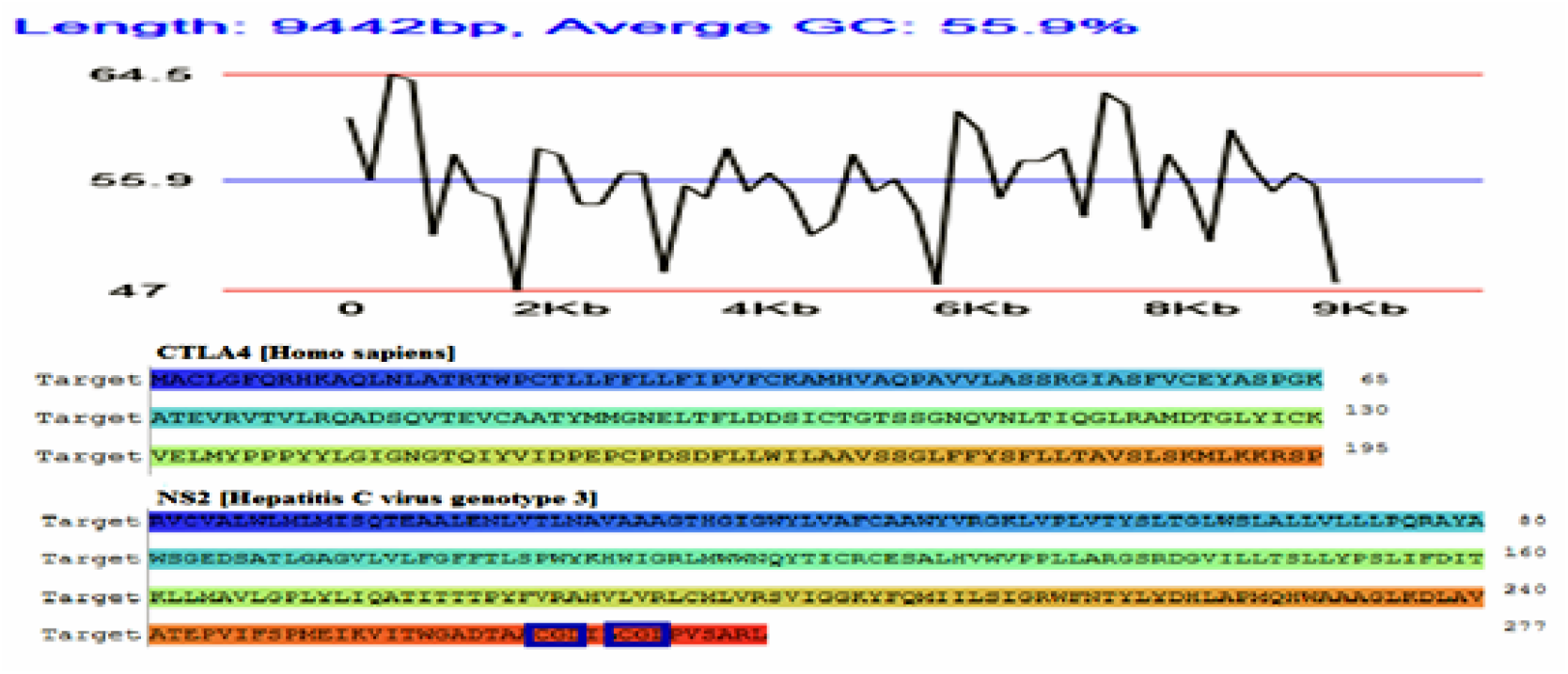
GC Content Distribution of HCV genotype 3.

To design of CTLA-4 primer was used Primer3 developed at NCBI using BLAST and global alignment algorithm to screen primers against selected database. The advanced and reverse primers were used to determine the methylation state of the CpGs within the CTLA-4 (Figure 9).

**Figure 9.**
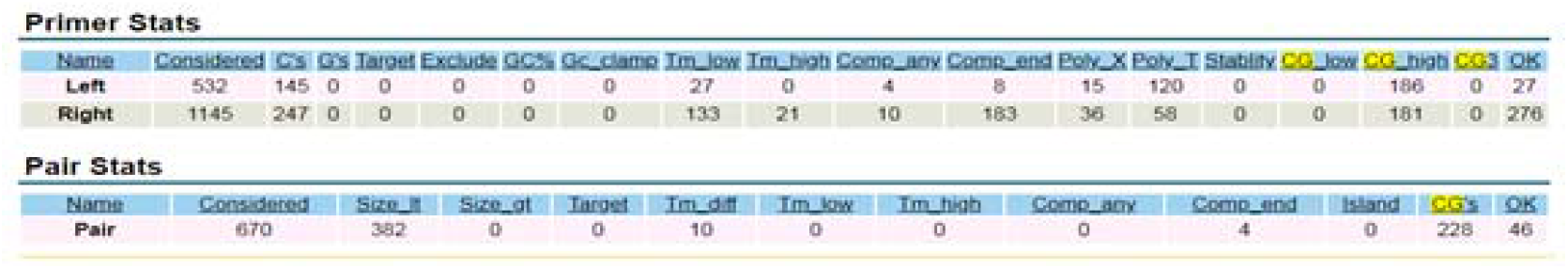
Primer nucleotide sequences of CTLA-4.

The DNA oligonucleotide sequence was altered by overlaying a C and G base on to A and T bases to accurately reflect the sequence of methylation in the CTLA-4 (Figure 10).

**Figure 10.**
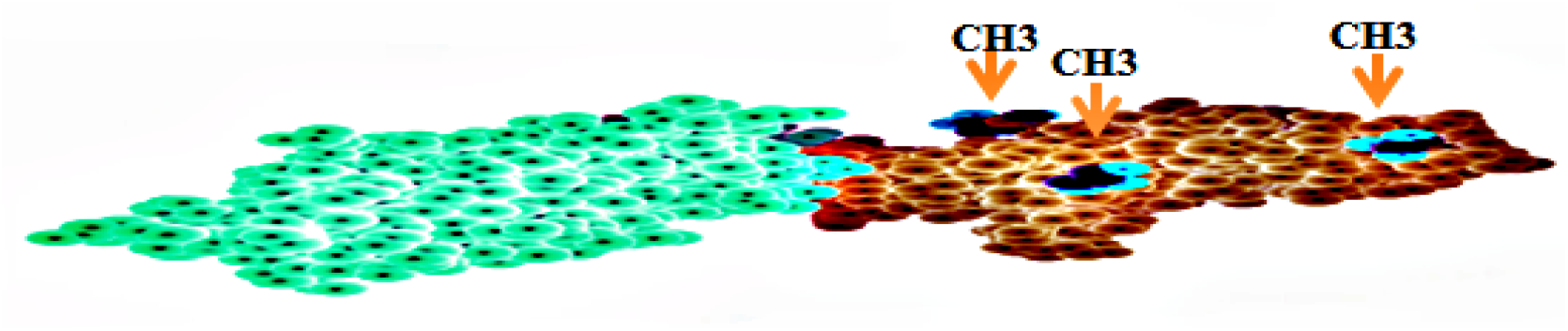
Methylation of CTLA-4.

## Discussion

Our *in silico* assessment suggests that epigenetic mechanisms may be underlying to triggering of autoimmune thyroiditis in individuals with HCV infection through of methylation of the CTLA-4 Gene promoter region.

Various extrahepatic disturbances are described associated to HCV infection, although the thyroids diseases be are generally the endocrinopathy most commonly diagnosed. The mechanisms of induced thyroid disease are complex but not entirely comprised (14).

Epigenetic disequilibrium promptly induces the progression of autoimmunity through regulation of immune cell functions (15). In diverse autoimmune diseases the epigenetic effect seems to perform a significant function in its triggering (16). Associated with this CTLA-4 that is essential immune checkpoint receptor regulating the T-cell activation, conferring predisposition to thyroid autoimmunity (17).

The human CTLA-4 Gene begins starting from 202 949·6 kb from the p-terminus of chromosome 2 and encompasses 6·2 kb on chromosome region 2q33 (18). NCBI based we present the location of structure of the CTLA-4, that has cytogenetic location in the 2q33.2, which is the long (q) arm of chromosome 2 at position 33.2, and molecular location in base pairs 203,867,771 to 203,873,965 on chromosome.

CpGi are regions with at GC frequency above 50%, at least 200 bp, and an observed-to-expected CpG ratio above 60%. CpGi describe potential CpGi regions employing the method reported by Gardiner-Garden and Frommer (19). We use the default parameters as proposed by Deaton and Bird (13) to identify the potential CpGi, and use as parameters length of DNA sequence > 300bp; CpG observed/CpG expected ratio > 60% and C+G% > 50%.

The cytosines in CpG dinucleotides are able to be methylated to form 5-methylcytosines, and circa of 70% to 80% of CpG cytosines are methylated in the mammals (20). In our analysis, the sequence alignment of CTLA-4 and HCV genome revealed the presence of two putative promoters and two CpGi in the 5’ genomic region of HCV genotype 1 Gene, one putative promoters and one CpGi in the 3’ genomic region of HCV genotype 2 Gene, and the presence of two putative promoters and two CpGi in the 5’ genomic region for HCV genotype 3 Gene. We did not find any published articles that associated the sequences alignment of CTLA-4 and HCV genotype 3 Gene, CTLA-4 and HCV genotype 2 Gene, and CTLA-4 and HCV genotype 1 Gene.

The Hashimoto’s thyroiditis and Grave’s disease are the most frequent autoimmune manifestations that occur in the thyroid (21). Recent studies demonstrated that DNA methylation is present in autoimmune thyroid disease determining a substantial epigenetic effect, being verified hypermethylated gene loci of CTLA-4 simultaneously to T cell receptor signaling (22). Our *in silico* analysis and the primers for quantitative DNA methylation analysis of thryroid peroxidase, and the sequence alignment of CTLA-4 and thyroid peroxidase not revealed the presence of putative promoters and none CpGi, but the advanced and reverse primers used to determine the methylation state of the CpGs within the CTLA-4 demonstrated that the DNA oligonucleotide sequence altered by overlaying a cytosine and guanine base on to adenine and thymine bases accurately reflected the sequence of methylation in the CTLA-4.

Recent studies proved that HCV infection as an important infectious triggering factor of autoimmune thyroiditis (23), and although the precise way of this association still is uncertain, one of potentials mechanism is related to epigenetic alterations as demonstrated in our study. Thus, epigenetics alterations have been to play a role important in the trigger of autoimmune thyroiditis.

## Conclusion

The role of CTLA-4 methylation in the trigger of autoimmune thyroid disease in HCV infection was *in sílico* evaluated in the present study, and the results showed that CTLA-4 methylation can be a factor that induces the trigger of autoimmnune thyroiditis. Nevertheless, in vitro and in vivo researches are needed to better clarify the exact implicit mechanism in deregulation of thyroid peroxidase activities in HCV infection by CTLA-4 methylation. We conclude that, in susceptible individuals, hypermethylation promotes reduced CTLA-4 expression and increases the risk of autoimmune thyroiditis in HCV-infected individuals.

